# Patient-derived glioblastoma cells (GBM) exhibit distinct biomechanical profiles associated with altered activity in the cytoskeleton regulatory pathway

**DOI:** 10.1101/2020.07.16.207233

**Authors:** Amelia Foss, Michele Zanoni, Woong Young So, Lisa Jenkins, Luigino Tosatto, Daniela Bartolini, Michael M Gottesman, Anna Tesei, Kandice Tanner

## Abstract

Glioblastoma multiforme (GBM) is the most commonly diagnosed brain cancer in adults, characterized by rapid proliferation and aggressive invasion into the stroma. Advances in our understanding of the molecular subtypes of GBM have provided attractive druggable targets. However, the high degree of heterogeneity both among patients and within individual tumors has proven a significant challenge for the development of effective therapies. We hypothesized that this heterogeneity is also represented in the mechanical phenotypes of GBM, as the physical properties of tumor tissue strongly influence elements of tumor progression including cell cycle regulation, migration, and therapeutic resistance. To assess these phenotypes, we employed optical trap-based active microrheology to determine the viscoelastic properties of patient-derived GBM cells in 3D hydrogels mimicking the brain ECM. We found that each GBM cell line had a distinct rheological profile as a function of treatment status, and cell lines could be further characterized by strong power law dependence describing intracellular viscoelastic behavior. Single-cell phenotyping according to power law dependence was able to identify subpopulations of cells within the treatment-resistant line. Finally, proteomic analysis indicated that altered mechanical profiles were associated with differential cytoskeletal regulation, particularly in actin - and myosin-binding pathways. This work suggests that evaluating mechanical properties may serve as a valuable strategy for the further stratification of these tumors, and encourages the investigation of cytoskeleton regulation as a potential therapeutic target for GBM.

## Introduction

Glioblastoma multiforme (GBM) is the most lethal cancer of the central nervous system, and most patients succumb to the disease within 1-2 years after diagnosis[1]. Transcriptomic and histopathological analysis have determined that GBM tumors adopt a multitude of phenotypic, cellular, genetic, and epigenetic subtypes[2, 3]. These molecular sub-types are characterized by amplification, absence, or mutation of genes that encode for growth factor receptors, metabolic regulators, and known tumor suppressors[2, 3]. This wealth of molecular data provides attractive and actionable biomarkers. However, clinical trials based on this stratification have not yielded curative treatment[1]. Moreover, recent data have shown that these sub-types are plastic, as an interconversion of distinct states is observed during tumor evolution[3]. It is thought that treatment efficacy is obfuscated due to the multifactorial diversity found intratumorally and across patients. Thus, further stratification of “multiformes” may be needed to develop effective therapeutic interventions.

GBM plasticity may in part be due to specific components of the brain microenvironment[4–7]. The physical properties of the brain microenvironment have been shown to influence drug response and tumor growth[4, 8, 9]. In addition, motile GBM cells can remodel the surrounding matrix, which, in turn, increases motility [4, 8–11]. This dynamic coupling between cell and environment also changes the phenotype of the cells themselves[11, 12]. The combined viscous and solid behavior that defines the single cell mechanical phenotype may also be a viable target to assess phenotypic plasticity. This mechano-phenotype regulates the kinetics and diversity of enzymatic processes which in turn drive cell fate[13–15]. Subcellular components such as the cytoskeleton, nucleus and a copious distribution of cytoplasmic proteins, enzymes and solutes drive the viscoelastic behavior of single cells[13–17]. Thus, defining these mechanical states may offer additional insight in what may drive the interchange between sub types in GBM. However, it is not known if single cells adopt distinct mechanical phenotypes as a function of subtype or within a given subtype for patient derived GBM cells.

Real-time mechanical mapping of GBM tumors in patients has been achieved using modalities based on magnetic resonance[18–20]. Using single and multi-frequency magnetic resonance elastography, correlations between mechanical properties and tumor grade and mutational status have been reported[21]. The technique has also been successful in resolving the relative importance of the fluidity for distinct brain tumors[18]. Intratumoral heterogeneity can be distinguished for regions corresponding to ~100s of cells. However, single cell resolution has not been demonstrated at this time *in vivo.* One strategy to address this knowledge gap would be a combined approach where one employs the use of biomimetic platforms that mimic the *in vivo* GBM microenvironment with that of single cell rheological modalities such as brillouin microscopy, and optical and magnetic tweezers. Using optical tweezer based active microrheology, we have previously shown that we can map the mechanical coupling of tumor cells embedded in distinct 3D extracellular matrix (ECM) hydrogels[16]. Here, we aimed to determine if patient derived GBM cells display heterogeneities in their mechanical phenotype similar to the molecular heterogeneity observed in patients.

We reasoned that the intratumor heterogeneity in clinical presentation and genetic characteristics also includes heterogeneity in mechanical phenotype and cytoskeletal dynamics, which may contribute to differing tumor progression and altered invasion profiles. To investigate this question, we embedded patient-derived GBM cells into 3D hyaluronic acid-based hydrogels to mimic the extracellular matrix environment of the brain. We determined that among three patient-derived GBM cell lines and a model cell line, each line had a distinct viscoelastic profile indicating cell stiffness and hysteresivity, or how liquid- or solid-like a substance behaves. Moreover, we determined that within each cell line, a range of mechanical phenotypes is observed. However, classification of cells based on power law dependence revealed that a cell line obtained from a relapsed patient stratified into two distinct populations, whereas the treatment naïve cell lines showed single populations with varying widths of distribution. Immunofluorescence shows altered patterns of actin cytoskeletal organization among lines. Supporting these data, cytoskeletal regulation pathways, including cytoskeletal protein-binding and myosin-binding pathways are significantly altered among cell lines according to mass spectrometry analysis. Specifically, Myosin IIC, associated with cytoskeleton dynamics, cell adhesion, and tumor cell invasion, is differentially abundant among lines, and serves as a promising target for further investigation.

## Results

### Mapping the intracellular heterogeneity of a single cell in 3D hyaluronic acid

In optical trap (OT)-based rheology, local piconewton forces are applied to objects such as beads and organelles [22–24]. From the induced displacement of these objects, determination of the underlying local physical properties of the material can be achieved, with the caveat that the mesh size of the material is smaller than the probe. Thus, we first asked if we can map cytoplasmic heterogeneity using 1 μm-diameter polystyrene beads as local sensors. To address this question, we employed a biomimetic model of the brain ECM microenvironment using the immortalized U87 GBM cell line. Cells were harvested and incubated in media supplemented with 1 μm-diameter polystyrene beads. These cells were embedded in 3D hyaluronic acid hydrogels at densities that minimized nearest neighbor effects and ensured cells saw an isotropic distribution of matrix (Figure 1A). OT-based active microrheology was performed on intracellular beads with a total amplitude of oscillation of 200 nm (20nm per frequency (ω)) at a power of 200mW at the back aperture over a range of frequencies spanning 7-15 kHz (Figure 1B). Using vis a vis calibration and FDT, we quantitated the complex modulus, G* for each bead. G* = |G*|exp(iδ) = G’ + iG”, can be then broken down into magnitude |G*| = (G’^2^ + G”^2^)^1/2^, in which G’ represents the storage modulus and G” represents the loss modulus. We also assessed crossover frequency the two components, which indicates when the material properties transition from solid-like to liquid-like. We determined the local physical properties for three to five intracellular beads randomly distributed in the cytoplasm of individual cells (Figure 1C). We found that, for a given cell, the values of G’ and G” for each bead varied from 6 Pa – 2 kPa for ω ranging from 7-15kHz. For each bead, we observed a strong power law dependence for frequencies greater than 400Hz (HF) compared to a relatively shallow curve at frequencies below 400Hz (LF).

**Figure 1.**
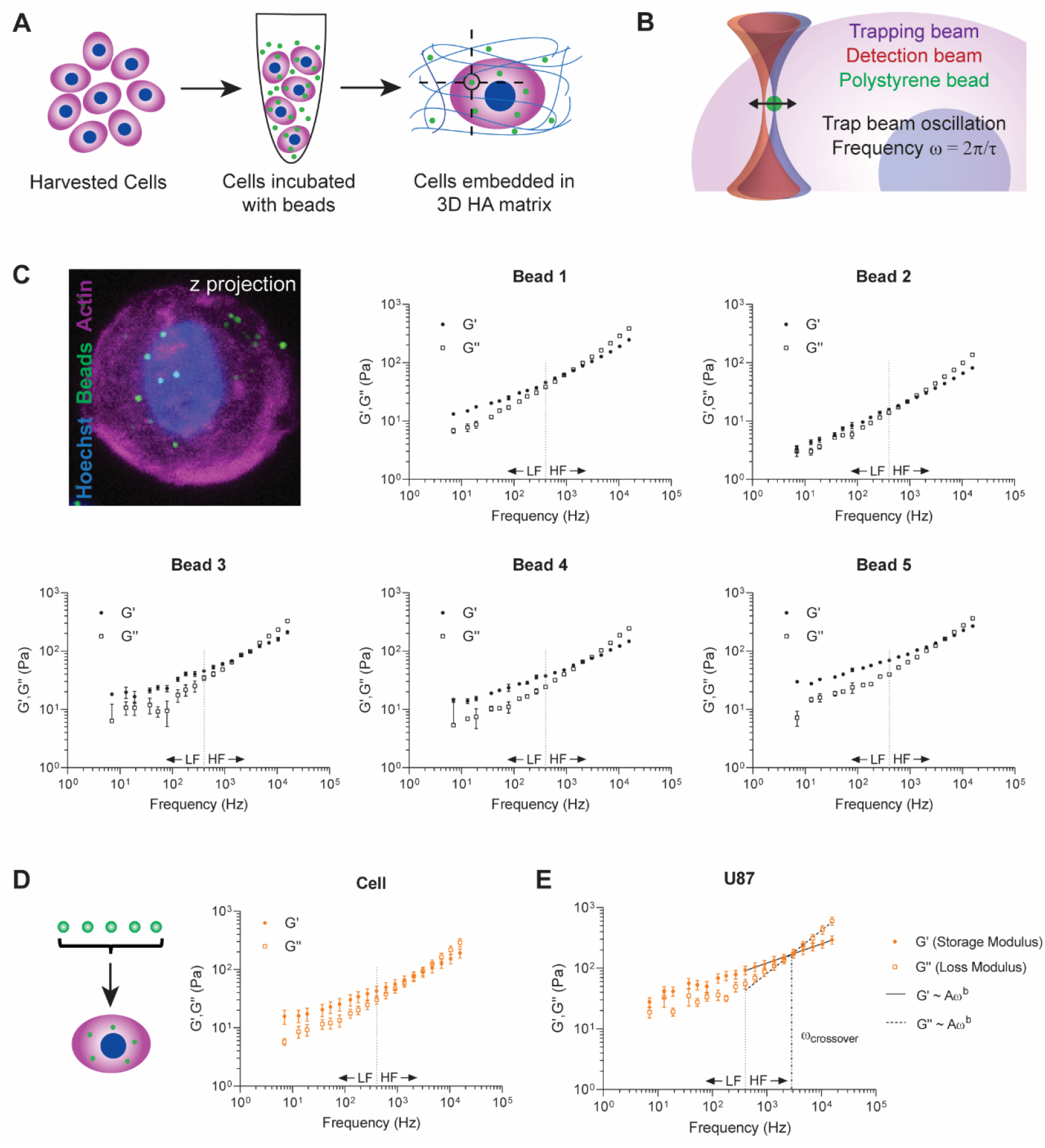
Optical trap-based microrheology maps intracellular mechanic heterogeneity of GBM cells in biomimetic model of brain ECM. **A)** Schematic of experimental design. U87 cells were harvested and incubated with 1 μm polystyrene beads in culture medium such that cells internalized the beads. Cells were then embedded in hyaluronic acid hydrogels and incubated overnight. Measurements of individual beads within cells were conducted 20-28h later. **B)** Schematic of optical trap measurement. Trap beam oscillation frequencies ranged from 7Hz-15kHz, and each bead underwent 7 measurements at each frequency. **C)** Micrograph of cell with internalized beads and G’,G” values for 5 individual beads in a single representative U87 cell (mean ± s.e., n ≤ 7 measurements). Micrograph staining includes Hoechst (nucleus, blue), phalloidin (actin, magenta), and polystyrene beads (green). **D)** Schematic and rheological values for a single cell (mean ± s.e., n = 5 beads). **E)** Average G’,G” values for U87 cell line (mean ± s.e., n = 30 cells). Low frequencies (LF) < 400 Hz. High frequencies (HF) >400 Hz. HF values fit to power law G’,G” ~ Aω^b^. For U87 cells, power law coefficients A, exponents b, and 95% confidence intervals are as follows. U87 G’: A = 13.71 (11.99, 15.43), b = 0.316 (0.3015, 0.3305); U87 G”: A = 0.6091 (0.3175, 0.9007), b = 0.7095 (0.6573, 0.7617). The ω_crossover_ is the point at which the G’ and G” power law curves intersect. For U87 cells, ω_crossover_ = 2.73 kHz.

We then quantified the average mechanical properties for a given population of cells. First, the phenotype, G’, G”, of each cell was determined by averaging measurements for each bead within a given cell (Figure 1D). We then averaged the values for thirty cells. We observed G’ vs G” values of 18.8 – 602.5 Pa. and a crossover frequency at ω = 2.73 kHz. The distinct strong power law dependence following G’, G” ~ A(ω)^b^ at ω > 400Hz (HF) and shallow slope for frequencies below 400Hz (LF) is maintained for an ensemble population of cells (Figure 1D).

### Patient derived GBM samples show distinct mechanical properties as a function of treatment status

We next examined the rheological properties of primary GBM cells. We chose three distinct cell lines: two that were treatment naïve and one that was derived from a relapsed tumor following radiation and chemotherapy (Figure 2A). We confirmed that the intracellular heterogeneity observed in the immortalized cell line was also present in primary cells (Supplemental Figure 1A-C). For each primary cell line, the mechanical phenotype is dominated by the elastic modulus where the crossover frequency ranged from 2.5-4kHz (Figure 2B). As observed with the immortalized cell line, values of G’ and G” obeyed a power law at high frequencies in the primary lines. For the treatment naïve cell lines the value of exponent b ranged from 0.231-0.259 for G’ and from 0.685-0.712 for G”. Similarly, for the treatment resistant line, G’ followed a weaker power law dependence with a value of 0.268 for the exponent b, and a stronger dependence for G” where the value of the exponent b is 0.688. The treatment-resistant line showed a markedly higher ω_crossover_ at 4.06 kHz compared to the lower crossover frequencies of treatment-naïve lines (ω_crossover_ = 2.94 kHz, 2.95 kHz for GB40 and GB70, respectively) (Figure 2B, Supplemental Figure 2B). We can then transform these data into plots of complex moduli. These plots show that the absolute value of G* is dominated by the storage modulus whereas the component from the loss modulus dominated the exponent of the power law dependence (Figure 2C). To assess relative stiffnesses, we compared the Log_2_ ratios of G*(ω) for each cell line. From this analysis, one naïve cell line (GB70) was stiffer than the primary and immortalized cell lines (GB34 and GB40, and U87, respectively) (Figure 2D). The immortalized cell line showed a similar stiffness compared to treatment resistant cell line at high frequencies. As predicted, at the higher frequencies, G* showed a stronger power law dependence where slopes ranged from 0.47 to 0.56 (Figure 2C)[25]. To assign a mechanical phenotype to each of the cell lines, we examined the power law dependence of G* ∝ Aω^b^ for each population, where materials that behave like semi-flexible polymers follow b=0.75, with b=0.5 indicative of flexible polymers. We determined that treatment resistant cells behaved relatively more like flexible polymers compared to treatment naïve cells, with a b=0.47 for the GB34 line compared to b = 0.48 and b = 0.49 for GB40 and GB70 lines, respectively.

**Figure 2.**
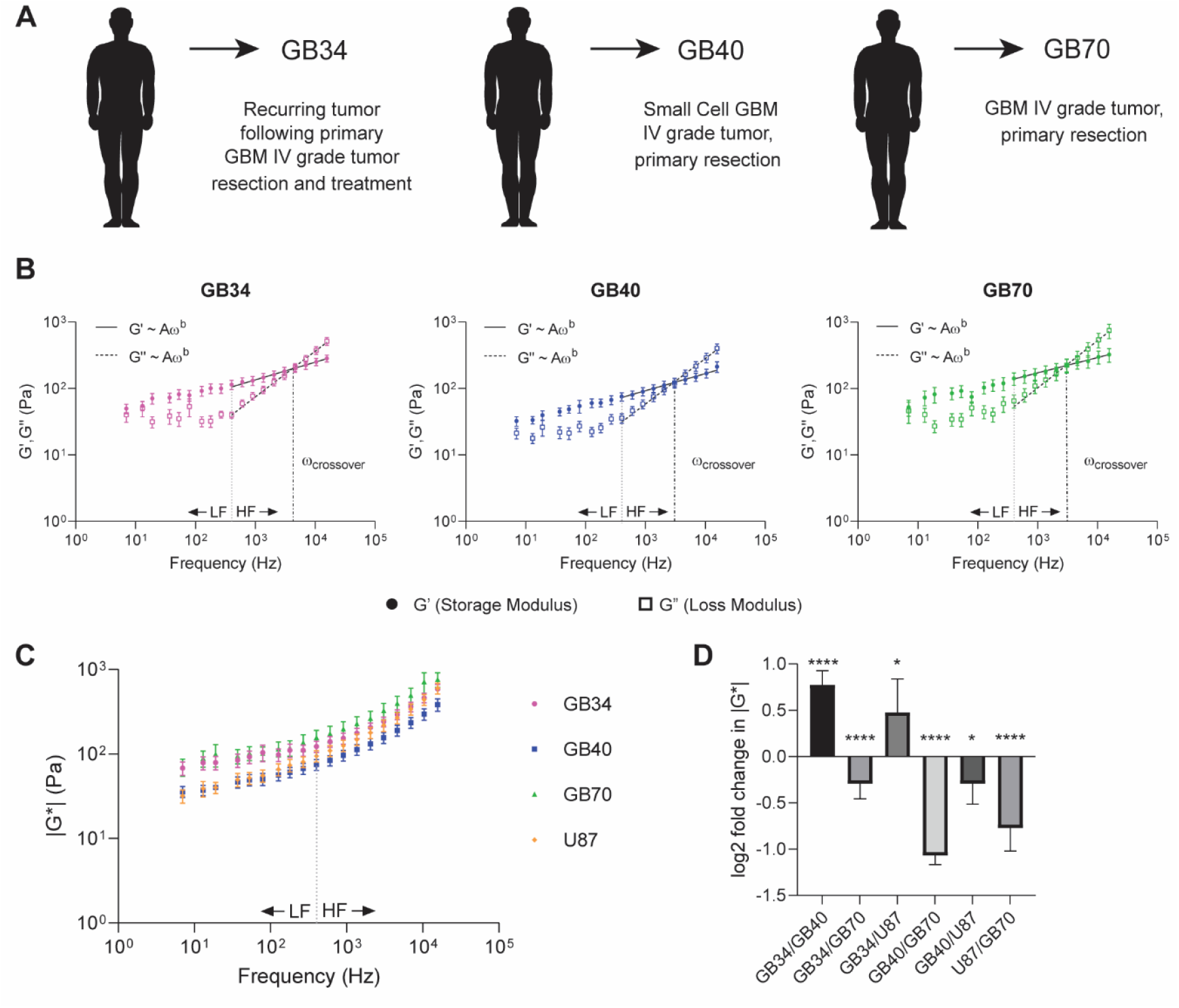
Patient-derived GBM cell lines show distinct rheological profiles. **A)** Three primary GBM cell lines were derived from patients. The patient providing GB34 cells underwent standard resection and treatment, and the cells are derived from a biopsy of the recurring tumor. GB40 and GB70 cells are treatment-naïve. **B)** Average G’, G” values for each primary GBM cell line (mean ± s.e., n ≥ 17 cells). Power law coefficients A, exponents b, and 95% confidence intervals are as follows. GB34 G’: A = 21.23 (18.18, 24.28), b = 0.2675 (0.2508, 0.2843); GB40 G’: A = 15.49 (12.73, 18.25), b = 0.2587 (0.2378, 0.2795); GB70 G’: A = 34.87 (29.88, 39.87), b = 0.2308 (0.2139, 0.2477). GB34 G”: A = 0.6455 (0.4317, 0.8593), b = 0.6879 (0.6517, 0.724); GB40 G”: A = 0.5129 (0.3234, 0.7025), b = 0.6854 (0.645, 0.7257); GB70 G”: A = 0.749 (0.4735, 1.024), b = 0.7116 (0.6715, 0.7517). The crossover frequencies for each line are as follows: GB34 ω_crossover_ = 4.06 kHz; GB40 ω_crossover_ = 2.94 kHz; GB70 ω_crossover_ = 2.95 kHz. **C)** The magnitude of the complex viscoelastic modulus |G*(ω)| is plotted for primary and immortalized cell lines (mean ± s.e., n ≥ 17 cells). **D)** Pairwise log2 ratios of |G*(ω)| across frequencies comparing all lines. Significance was determined using a 2-way ANOVA (* p < 0.05, ** p < 0.01, *** p < 0.001, **** p < 0.0001).

### Single cell phenotyping using power law dependence identifies sub-populations in treatment resistant GBM cells

GBM tumors are comprised of cells that show a diversity of phenotypes hence the use of “multiforme” in its tumor classification[1, 3]. Thus, we revisited our individual cell measurements to understand the distribution of the mechanical phenotypes. Among all cell lines, the individual cell measurements ranged from ~10 – 2500Pa for the full spectrum of frequencies, displaying significant heterogeneity within each line (Figure 3A). Having observed a robust power law at higher frequencies, we then calculated the individual exponents on a single cell basis (Figure 3B). We observed that the treatment resistant GB34 cells segregate into two populations, cells that behave like flexible and semi-flexible polymers. In contrast, the treatment naive cell lines show single populations, one (GB40) that can be classified with phenotypes akin to flexible polymers whereas another cell line (GB70) showed a broad distribution of states. In contrast, phenotypes of the immortalized cell line, U87 clustered more closely with phenotypes akin to semi-flexible polymers. We also observed these population dynamics in single cell analysis of intracellular hysteresivity, as resistant GB34 cells appear to segregate into two populations, GB70 cells show a broad distribution of characteristics, and the remaining treatment naïve lines cluster close together (Supplemental Figure 2A).

**Figure 3.**
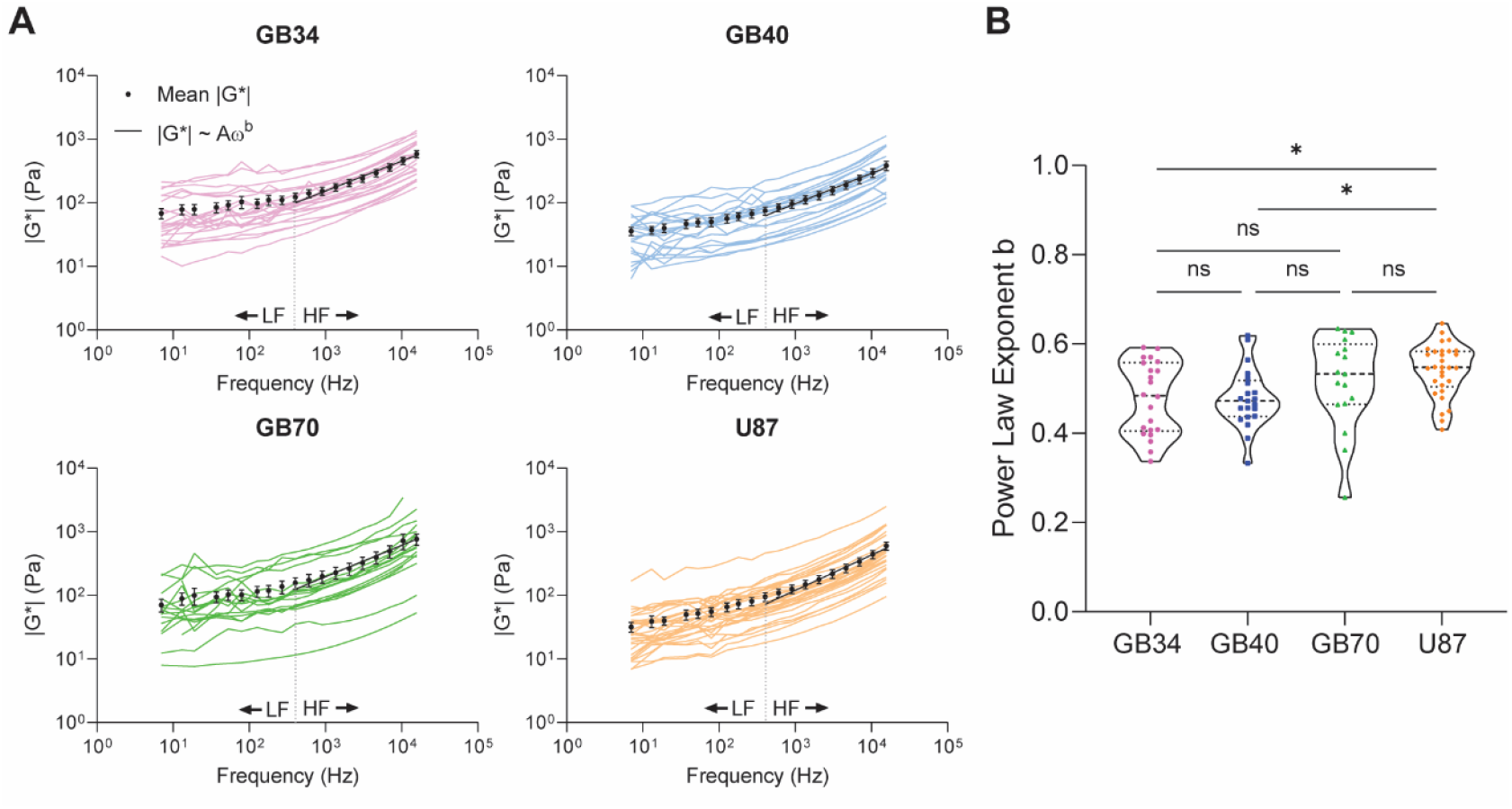
Single cell analysis demonstrates line-specific power law-dependent rheological profiles. **A)** Mean |G*(ω)| values for each cell within its respective population accompanied by population average (mean ± s.e.). HF population mean values were fit to the power law G* ~ Aω ^b^. The power law coefficients A, exponents b, and 95% confidence intervals are as follows: GB34 |G*|: A = 5.897 (3.142, 8.652), b = 0.471 (0.4186, 0.5234); GB40 |G*|: A = 3.545 (1.927, 5.163), b = 0.4782 (0.4271,0.5293); GB70 |G*|: A = 6.431 (1.854, 11.01), b = 0.4947 (0.4152, 0.5742); U87 |G*|: A = 2.593 (0.979, 4.206), b = 0.555 (0.4861, 0.624). **B)** The HF values of individual cells were fit to the power law G* ~ Aω^b^. The exponent b for each cell is displayed, along with median and first and third quartiles for each line. Significance was determined using a one-way ANOVA with Tukey’s multiple comparisons test. (* adjusted p value < 0.05)

### GBM lines show distinct patterns of cytoskeletal organization

Mechanical phenotypes can be modulated by many factors such as treatment and culture conditions (i.e. primary vs immortalized). Thus, to assess what molecular mechanisms may regulate the differences in mechanical phenotypes, we focused on two of the treatment naïve cell lines. Differential analysis of the proteome revealed that Myosin IIB (MYH10) and IIC (MYH14) were upregulated in the GB40 cell line compared to the more heterogenous GB70 line (Figure 4A-B). Myosin IIB and IIC have been implicated as regulators of actin organization [26, 27]. In cells, actin-dependent regulation of cytoskeletal architecture provides a significant contribution to single cell mechanical properties. Thus, we assessed actin structures using immunostaining (Figure 4C). In the cell line that showed a narrow distribution akin to flexible polymers, we observed actin structures assembled into multi-foci within the cytoplasm with minimal cortical staining. However, in the second cell line that showed broad distribution of mechanical states, two distinct of actin structures and distributions were observed: 1) multi foci and 2) diffuse with strong cortical staining. Shape analysis of these cell lines also showed a difference in circularity, where GB40 cells with stronger punctate staining were less spherical than the GB70 cells (Supplemental Figure 3). Furthermore, pathway analysis indicates significant upregulation of cytoskeletal organization functions in the GB40 line, including actin and myosin binding (Figure 4D).

**Figure 4.**
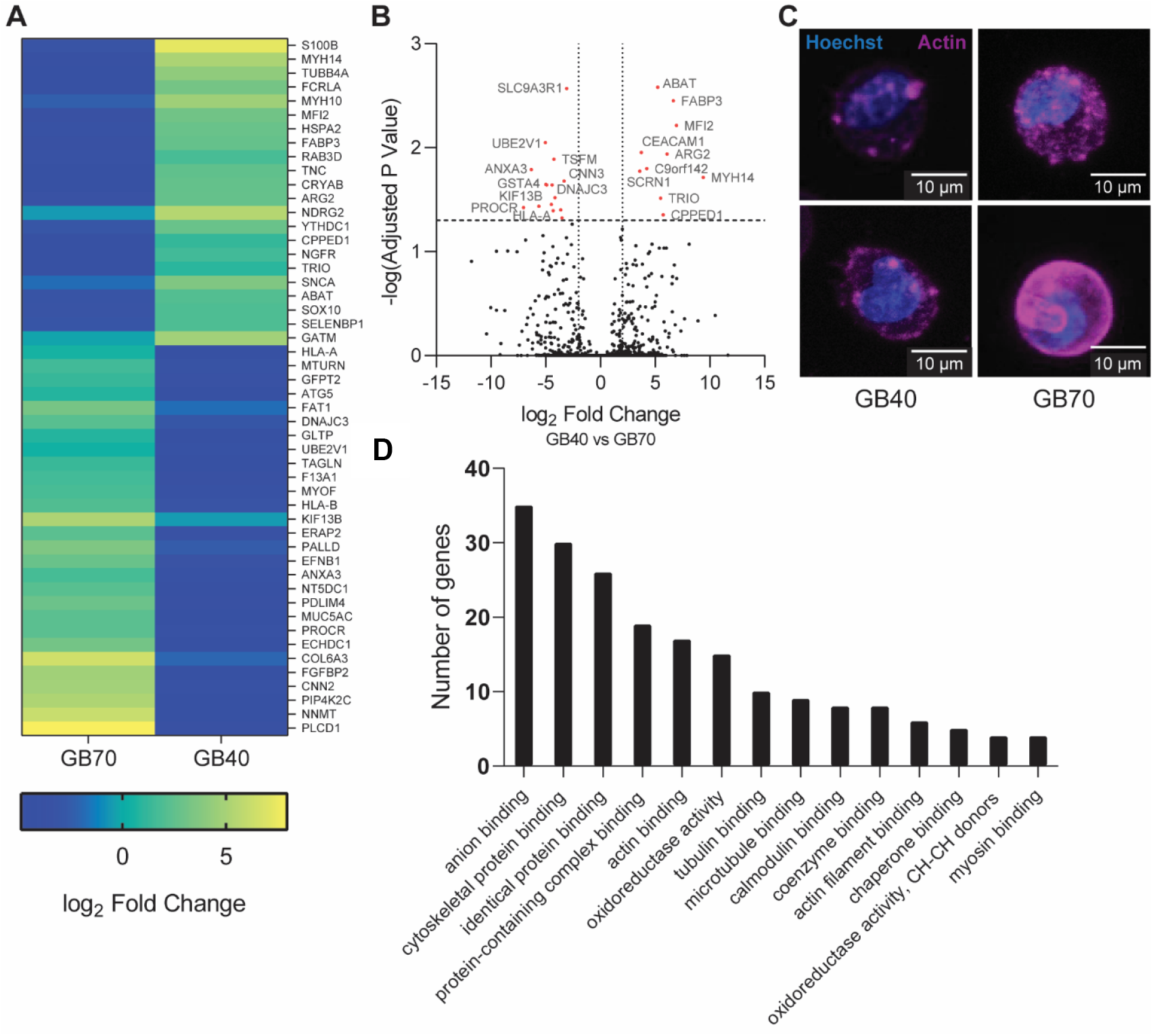
Primary GBM lines show differential abundance of proteins associated with cytoskeletal organization. **A)** Mass spectrometry analysis indicates differentially abundant proteins between GB40 and GB70 lines. Heatmap displays top 50 statistically significant (p < 0.05) most differentially abundant proteins. Fold change was determined relative to the global median log2 values across all samples. **B)** Volcano plot displaying log_2_ fold change vs the negative logarithm of the adjusted p value, calculated using the Holm-Sidak method with α < 0.05. Horizontal line indicates the negative log of adjusted p value with α = 0.05. Vertical lines indicate a log_2_ fold change of ± 2. **C)** Representative micrographs of cells from GB40 and GB70 lines. Staining includes Hoechst (nuclei, blue) and phalloidin (actin, magenta). **D)** Pathways enriched in the GB40 line compared to the GB70 line among top 150 most statistically significant differentially abundant proteins. Pathway analysis was limited to Gene Ontology (GO) Function terms with FDR < 0.05.

## Discussion

In spite of our knowledge of molecular sub-types of GBM, curable treatment remains elusive[1]. Treatments include chemotherapy (temozolomide), radiation, and surgical resection, and occur only if these interventions do not impair cognition and quality of life [1,28]. Phenotypic plasticity and microenvironmental elements are potent factors affecting drug sensitivity and efficacy [1, 3]. These factors are further complicated by intratumor and intertumoral heterogeneities in patients, which have thus far been challenging to model at high resolution [1–3, 7, 29]. Our lab has previously measured the microscale mechanical coupling of tumor cells with that of a 3D biomimetic hydrogel [16]. Here, in an effort to understand mechanical heterogeneity in glioblastoma, we employed optical trap based active microrheology to characterize single cell mechanical phenotypes of patient derived primary GBM cells in a 3D brain mimicking matrix. We measured the single cell mechanophenotype of three patient-derived GBM primary cell lines and an immortalized cell line. Each of these had a distinct viscoelastic profile indicating cell stiffness and hysteresivity, or how liquid- or solid-like a substance behaves. Examination of power law dependence at higher frequencies revealed that the treatment resistant GBM cells can be grouped into two distinct populations, while most of the treatment naïve cell lines were centered on a single population and the other showed a broad distribution of states. For this cell line we determined that these cells adopted multiple states with different patterns of actin cytoskeletal organization.

The mapping of cellular mechanics has been achieved using multiple modalities including optical and magnetic tweezer based active rheology, passive rheology, atomic force microscopy and Brillouin microscopy[17, 30, 31]. Measurements of single cells in suspension can be performed using an optical stretcher and the high throughput, RT-DC system[32]. Previously, intracellular measurements obtained for yeast cells and normal mammalian cells ranging from neurons and fibroblasts to breast epithelial cells have been achieved using AFM and Brillouin microscopy [33–35]. These values ranged from ~ 0.5-2kPa for Young’s modulus and a calibrated longitudinal modulus respectively for a relatively narrow frequency range and non-overlapping regime[33–35], and interestingly, single cell measurements revealed a difference in mechanical properties due to differentiation status in fetal neural progenitor cells[14]. Here, we determined that the GBM cells are comparatively softer, where the measured complex moduli ranged from 0.02-1 kPa across our multiplexed 20 frequencies. This is in line with previous measurements using OT based microrheology of 2 micron beads in breast cancer cells in cultured on 2D micropatterned substrates and 1 micron beads of breast cancer cells cultured in 3D hydrogels[16, 36]. Moreover, our cells showed differences in stiffnesses due to treatment status, where treatment naïve cells were softer than the treatment resistant. This result is consistent with changes of mechanical phenotypes due to cell fates observed in comparison of differentiation status and drug resistance[14, 33]. This discrepancy between the OT measurements and the non-probe-based techniques can be due to four factors. First, mechanical phenotype is strongly dependent on the length and temporal scales of the technique. Second, mechanical properties obtained using probe-based techniques such as OT/MT and particle based passive rheology are highly dependent on probe size. Third, healthy, normally-programed cells are often “stiffer” than their cancerous counterparts. Last, mechanical properties are dependent on culture conditions, such as adherent or suspended culture, and dimension (2D vs 3D)[9, 37–39].

Access to broad frequencies allowed us to examine a dynamic range of complex behaviors. Our measurements compare well with previous measurements, where G* is characterized by a low frequency elastic plateau regime and a high frequency liquid-like regime characterized by powerlaw frequency dependence[25, 38, 40]. In particular, actin cytoskeleton dynamics dominate the mechanical response due to the size of the probe used in these experiments. At higher frequencies (>400Hz), the exponent from the formulaic power dependence is equal to 1 for purely viscous response. If contributions due to the cytoskeleton are the largest contributors, then these values are 0.5, 0.75 and 0.875 for transverse and longitudinal modes of relaxation respectively [41]. We previously demonstrated that this exponent can vary from 0.2-0.8 for breast cells cultured in 3D hydrogels based on the chemistry of the hydrogel and perturbation of the cytoskeleton[16]. Similar examination was performed for cells cultured in 2D using high frequency AFM[40]. In these measurements, a double exponential fit was used for the elastic and liquid like regime. This value ranged from 0.15 – 0.9 based on cell type and pharmacological inhibition of actin. In a similar comparison, an exponent of 0.41 was calculated for a breast cancer cell line[42]. Here, our measurements showed a range from 0.47-0.56.

We used this exponent to categorize the single cells in an effort to describe a complementary mechanical phenotype reminiscent of the molecular phenotypic classification used for GBM. In this work, we observed a difference due to treatment status. It remains to be seen, however, if the mechanical phenotype can be linked to mutation status as observed for MRE measurements[21]. Instead our data hint at the importance of factors that direct cellular mechanics, such as the actomyosin pathway, as potential regulators of the “multiformes” within the GBM population.

Non-muscle myosin proteins have been implicated in tumor proliferation and invasion. Briefly, decreased activity of Myosin IIA and IIB in GBM cells, particularly Myosin IIA, resulted in reduced tumor cell invasion *in vitro* and in a rodent model [43–45]. However, decreased Myosin IIA expression also enhanced proliferation in a manner dependent on the mechanical properties of the tissue microenvironment[45]. Our data, obtained using direct comparison of the proteome of two representative treatment naïve cell lines, also revealed differential expression of cytoskeletal regulation pathways, including cytoskeletal protein-binding and myosin-binding pathways. Specifically, Myosin IIC, associated with cytoskeleton dynamics, cell adhesion, and tumor cell invasion within brain ECM, was differentially abundant among lines [26, 27, 46]. Our data presents additional evidence that these myosin isoforms may be key factors in GBM and serve as promising targets for further investigation.

## ACKNOWLEDGEMENTS

This research was supported by the Intramural Research Program of the National Institutes of Health, the National Cancer Institute.

## Methods

### Patient-derived glioblastoma cells

Primary human glioblastoma cells were obtained directly from patient tumor specimens processed within 2 hours of surgery, as previously described[47]. Briefly, adult GBM tissues obtained from patients underwent surgical resection and sampled by a pathologist were dissected and digested in an enzyme mix solution. Patient-derived GBM cells were plated in NeuroCult NS-A serum-free medium (StemCell Technologies, Vancouver, BC, Canada) supplemented with 20 ng/ml of epidermal- and 10 ng/ml of fibroblast-growth factor (Sigma-Aldrich), 1% penicillin/streptomycin (GE Healthcare, Milan, Italy) and 2% amphotericin B (Euroclone) and maintained under hypoxic conditions (1% O_2_) in a humidified 37 °C, 5% CO_2_ incubator. All the primary GBM cell lines were checked periodically for mycoplasma contamination using the MycoAlertTM Mycoplasma Detection Kit (Lonza, Basel, Switzerland). All Patient-derived biological samples were collected according to a protocol approved by Italian Local Ethics Committee (CE IRST IRCCS-AVR, protocol 2439/2018) and all the patients enrolled have signed an informed consent for the genetic analyses and for the use of the results for research purposes.

### Cell Culture

U87 cells were grown and maintained as previously described at 37°C in normoxic conditions (20% O_2_, 5% CO_2_) in Dulbecco’s Modified Eagle Medium (DMEM, 10% FBS, 1% Penicillin/Streptomycin, 1% L-glutamate)[48]. Patient-derived glioblastoma (GBM) cells (GB34, GB40, GB70) were grown and maintained at 37°C in hypoxic conditions (3% O_2_, 5% CO_2_) in NeuroCult™ NS-A Proliferation medium (STEMCELL Technologies, 05751) supplemented with 1% Heparin Solution (STEMCELL Technologies, 07980), 2% Amphotericin B (Fungizone, EuroClone, ECM0009D), 10ng/mL bFGF (Sigma-Aldrich, FO_2_91), 20ng/mL EGF (Sigma-Aldrich, E9644), and 1% Penicillin/Streptomycin. Cells were passaged when 70-80% confluent, and media were changed regularly.

### Sample Preparation

Prior to preparation, U87 cells were detached with 10mM EDTA for 15min at 37°C. GBM cells were mechanically detached or collected from suspension in growth media. Cells were pelleted, counted, and resuspended to a concentration of 3×10^6^ cells/mL. Polystyrene beads were added to the cell suspensions in a 1:20 ratio (1×10^10^ stock solution, Life Technologies, FluoSpheres™ Polystyrene Microspheres, F1308), and incubated with gentle mixing for 30-45 min at 37°C. Hyaluronic acid (HA) hydrogels were prepared concurrent with cell preparation. Frozen lyophilized thiol-modified hyaluronan (Glycosil^®^, Advanced BioMatrix, GS222) and polyethylene glycol diacrylate (Extralink-Lite^®^, Advanced BioMatrix, GS3008) were thawed and dissolved with degassed deionized water to 50mg/mL and 5mg/mL, respectively, and vortexed for 45 min. For each sample, a volume of 40μL of cell-bead solution was added to 160μL Glycosil^®^ and mixed thoroughly with a micropipette. Immediately thereafter, 40μL Extralink-Lite^®^ solution was added and thoroughly mixed, such that the final concentration of the hyaluronan was 33.3mg/mL. For optical trap measurements, samples were prepared by pipetting 33μL solution onto a glass microscope slide. A chamber was created by adding a coverslip to each side of the slide and across the top, fixed with double-sided tape. For immunofluorescence imaging, the gel solution was pipetted into plastic wells affixed to a microscope slide. The gels were allowed to polymerize for 1 hr. at 37°C, after which cell growth media was added to hydrate the samples. Polymerization continued overnight, and samples were measured or fixed at 24-28 h post preparation.

### Immunofluorescence Imaging

Samples were fixed in 4% PFA for 2 h at room temperature. Samples were washed 3 x 10 min with PBS, and blocked and permeabilized for 1 h at room temperature in PBST (PBS containing 0.1% Tween 20) + 0.5% Triton X-100. Samples were washed 3 x 10min with PBS, then incubated for 2 h at room temperature in a staining solution of PBST containing a 1:50 dilution of phalloidin and 1:1000 dilution of Hoechst 33342. Samples were washed with PBS 3 x 10 min prior to imaging. Samples were imaged on a Zeiss 780 laser scanning confocal microscope with Plan-Apochromat 40x/1.4 NA Oil DC M27 objective. Presented images are maximum intensity projections of 1 μm z stacks.

### Quantification of Circularity

Analysis was performed in FIJI using the built-in circularity plugin within the *Analyze Particles* command. Maximum intensity projections of the phalloidin channel of single cells (GB40 n = 9; GB70 n = 6) were thresholded and circularity was determined on a scale of 0-1 using the formula *circularity = 4π(area/perimeter^2^).* Differences between cell lines were determined using an unpaired, two-tailed Student’s *t* test.

### Mass Spectrometry

GBM cells (GB40 and GB70) were seeded at concentrations of 250,000 per well in a 6 well plate and starved for 24 hrs replacing conventional media with growth factor free media. Cells were then washed twice with PBS 1X and lysed in 50 μL of Mammalian Protein Extraction lysis buffer (Thermo Fisher Scientific) supplemented with phosphatase inhibitor cocktails 2 and 3 (Sigma Aldrich) and protease inhibitor (Sigma Aldrich) at 1:100. After 1 hour of incubation on ice, proteins were collected from supernatants after spinning down for 15 min at 12,000 x g. Collected supernatants were stored in 1.5 mL tubes at −80 degrees Celsius.

The filter-aided sample preparation (FASP) protocol [49] was used with minor modifications for the digestion of cell lysates. Briefly, lysates were reduced with 10mM DTT at 55°C for 30min, then diluted in a Microcon YM-10 filter unit with 8 M urea in 100mM Tris-HCl (pH 8.5 [UA]) and centrifuged at 20°C for 30 minutes at 14,000 x g. Samples were washed with 200 μL UA, and protein was alkylated using 50mM in UA for 6 minutes at 25°C. This was followed by centrifugation at 20°C for 30min at 14,000 x g, and excess reagent was removed. Samples were washed with 3 x 100 μL of 8 M urea in 100 mM Tris-HCl (pH 8.0 [UB]), then diluted to 1 M urea with 100 mM Tris-HCl (pH 8.0). Proteins were digested with trypsin at a ratio of 1:100 trypsin to protein w/w overnight at 37°C. Recovery of tryptic peptides was performed by centrifugation at 20°C for 30 minutes at 14,000 x g, then washing the filter with 50 μL 0.5 M NaCl. Peptides were desalted and acidified using a C18 SepPak cartridge (Waters, Milford, MA) and dried using vacuum concentration (Labconco, Kansas City, MO). Dried peptides were fractionated by high pH reversed-phase spin columns (Thermo Fisher Scientific), lyophilized, and solubilized in 4% acetonitrile and 0.5% formic acid in water. Fractions were separated on a 75 μm x 15 cm, 2 μm Acclaim™ PepMap™ reverse phase column (Thermo Fisher Scientific) with an UltiMate™ 3000 RSLCnano HPLC (Thermo Fisher Scientific) at a flow rate of 300 nL/min. Online analysis was performed by tandem mass spectrometry using an Orbitrap Fusion™ mass spectrometer (Thermo Fisher Scientific). Peptides were eluted into the spectrometer in linear gradients from 96% mobile phase A (0.1% formic acid in water) to 35% mobile phase B (0.1% formic acid in acetonitrile) over a period of 240 minutes. The Orbitrap mass analyzer was set to acquire data at 120,000 FWHM resolution, from which parent full-scan mass spectra were collected. Ions were isolated in the quadrupole mass filter, then fragmented in the HCD cell (normalized energy 32%, stepped ± 3%). Product ions were then analyzed in the ion trap.

Mass spectrometry data and label-free quantification were performed using the MaxQuant version 1.5.7.4 [50, 51]. Parameters were set as follows: variable modifications – methionine oxidation and N-acetylation of protein N-terminus; static modification – cysteine carbamidomethylation; first search performed using 20 ppm error; main search performed using 10 ppm error. Maximum of two missed cleavages; protein and peptide FDR threshold of 0.01; min unique peptides 1; match between runs; label-free quantification with minimal ratio count 2. The Uniprot human database from November 2016 (20,072 entries) was used to identify proteins, and statistical analysis was performed using Perseus version 1.5.6.0 [52]. Contaminants, reversed sequences, and proteins quantified in only one of three experimental replicates were removed. Label-free quantification values were base 2 logarithmized, and missing values were imputed from a normal distribution of the data. A two-way Student’s *t* test was performed, with a p-value cutoff of 0.05. Fold change was determined relative to global median base 2 logarithmic values across all samples. The adjusted p value and negative logarithm of the adjusted p value were calculated using the Holm-Sidak method with an alpha <0.05 in GraphPad Prism 8.0.2. Pathway enrichment was performed using the Cytoscape version 3.7.2, wherein the top 150 most statistically significant differentially regulated proteins were entered into the STRING Enrichment application. Output was limited to Gene Ontology (GO) Function terms with an FDR <0.05. Pathways with at least four proteins are plotted, with the exception of the parent terms “binding” and “protein binding” which contained 112 and 73 proteins, respectively. Of these 150 proteins, the fifty most differentially abundant proteins are plotted as a heatmap.

### Optical Trap Instrumentation and Setup

#### Instrumentation and Setup

Our home-built setup was used as previously described [22, 24]. The setup consists of a 1064 nm trapping beam (IPG Photonics, #YLR-20-1064-Y11) and a 975 nm detection beam (Lumics, #LU0975M00-1002F10D) mounted on 5-axis adjustable mounts (Newport, New Focus 9081).

The trapping beam is steered by a dual-axis acousto-optic deflector (AOD) (IntraAction, DTD274HD6), which oscillates the trap at frequency ω. Before entering the AOD, the trapping beam is linearly polarized by polarizing beam splitter cubes (Thorlabs, PBS23), and the beam is attenuated manually by half-wave plates (Thorlabs, WPH05M-1064) or electronically via analog output from a data acquisition (DAQ) card (National Instruments, PCIe-5871R FPGA). The AOD receives control signals from radio frequency generating cards (Analog Devices, #AD9854/PCBZ) with onboard temperature-controlled crystal oscillators (Anodyne Components, ZKG10A1N-60.000M), and the cards are controlled by the DAQ card. After passing through the AOD, an iris the doubly diffracted beam (i.e. 1^st^ order in both transverse axes). To detect the displacement of the trapping beam position, a beam sampler mirror (Thorlabs, BSF10-C) and neutral density (ND) filter (Thorlabs, NENIR210B) direct a small amount of power (~1%) onto the trap-assigned quadrant photodiode (QPD) (First Sensor, QP154-QHVSD). The trap beam is expanded by a lens pair (Thorlabs, LA1509-C, 100mm; AC508-200-B, 200mm) and directed through a dichroic mirror (Chroma, T1020LPXR).

The detection beam is expanded by a lens pair (Thorlabs, LA1131-C, 50mm; AC508-200-b, 200mm) and directed through the dichroic mirror, where it is coupled into and collocated with the trapping beam. A third telescopic lens pair (Thorlabs, LA113-C, 50mm; LA1384-CA, 125mm) expands both beams, and a dichroic filter cube (Chroma, ZT1064rdc-2p) sends both beams into the objective (Nikon, MRDO7602 CFIPLAN-APO VC60XA WI 1.2 NA), such that the trapping beam slightly overfills the back aperture of the objective. A high numerical aperture (NA), long working distance (WD) condenser (Nikon, WI 0.9 NA) collects light from the objective. A dichroic mirror (Chroma ZT1064RDC-2P) directs the detection beam through a relay lens that is positioned to image the back focal plane of the condenser onto the detection-assigned QPD. A bandpass filter (Chroma, ET980/20X) removes the trapping beam from the path to the detection-assigned QPD. Time-correlated ‘trap’ and ‘detection’ QPD signals are collected by analog inputs of the DAQ card.

A charge-coupled device (CCD) camera is mounted on the optical table such that camera position can be adjusted to a plane conjugate to the trapping beam AOD, back aperture of the condenser, and detection QPD. The Hx-nm constant relating the AOD RF control signal (in Hz) to the beam displacement (in nm) is calibrated by attenuating and focusing the beam on a coverslip and imaging the backscattered beam on the CCD camera.

The alignment of the beams and the back focal plane interferometer is confirmed before each experiment, and laser power is measured at the microscope backport with a power meter (Fieldmate, Coherent) and adjusted to 100 mW at the half-wave plate. A flow chamber is constructed from a microscope slide and coverslide with double-sided tape (Scotch) and loaded by capillary action with polystyrene beads (Life Technologies, F1308) suspended in water. To ensure proper alignment, a bead is trapped and the trap beam is oscillated while bead position is viewed in real time from the detection QPD signal, and the beam-coupling dichroic and QPD position are adjusted until oscillations in both transverse axes are centered on the QPD. A thermal power spectrum is recorded and fitted to a Lorentzian function to calculate the viscosity of the water to confirm system calibration. Before measurement, the camera pixel coordinates of the trap beam’s position are found by fitting a centroid to the intensity of an image collected of a trapped, stationary bead in water.

#### Sample Measurement

The condenser is placed in Kohler illumination and the sample is placed in focus. The bead of interest is positioned precisely in the center of the trap by scanning it through the detection beam in three dimensions using a piezo XYZ nanopositioning stage (Prior, #77011201) while voltages are recorded from the QPD. The characteristic V-nm relation, β, of QPD voltage vs. position was calibrated in situ by fitting the central linear region of the detector response to scanning the bead through the detection beam in the direction of the oscillations, giving β in V-nm, as previously described [24]. Once the bead is correctly positioned, the trap beam is oscillated while both trap and detection QPD signals are recorded. The oscillation is multiplexed as a superposition of sine waves of differing phase and frequency, with the same amplitude (25.4 nm) at each frequency. Frequencies are prime numbers, in the range of 3Hz-15kHz, to avoid interference. Four phases are interlaced to minimize the total amplitude of the composite waveform. The waveform is pulsed for 2 s, followed by 2s with the trap stationary to obtain active and passive spectra, respectively, and the sequence is repeated 7 times. Control and data collection were conducted in custom programs (National Instruments, LabVIEW).

#### Determination of Mechanical Properties

The optical trap stiffness is determined *in situ* from the active and passive power spectra using the active-passive calibration method [53, 54]. The linear viscoelastic mechanical response of the material is modeled by a generalized Langevin equation with additional force terms (a harmonic term accounting for the attractive potential of the optical trap and an acceleration memory term accounting for the elastic in-phase response) [54–57]. Mechanical response for undriven motion is described as 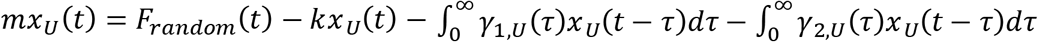, where t is time, *τ* is correlation time, *x_U_*(*t*), *x_U_(t)*, and *x_U_(t)* respectively the undriven bead position, velocity, and acceleration, *F_random_(t)* is the Brownian force, *k* is the optical trap stiffness, *m* is the bead mass, and *γ_1,U_(τ)* and *γ_2,U_(τ)* are respectively the real and imaginary parts of the undriven friction relaxation spectrum. Driven motion is described similarly as 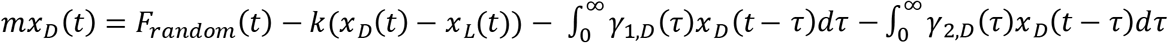, where *x_D_(t), x_D_(t)*, and *x_D_(t)* respectively the undriven bead position, velocity, and acceleration, *x_L_(t)* is the position of the optical trap, and *γ_1,D_(τ)* and *γ_2,D_(τ)* are respectively the real and imaginary parts of the driven friction relaxation spectrum. The fluctuation dissipation theorem identifies the undriven and driven friction relaxation spectra such that *γ_1, U_(τ)* = *γ_1,D_(τ)* and *γ_2, U_(τ)* = *γ_2,D_(τ)*, according to Onsager’s regression hypothesis.

The friction relaxation spectrum is related to the active power spectrum *R_L_(ω)* by the equation 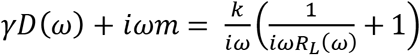 with probe frequency of the driving oscillation ω in rad•s^-1^. The optical trap stiffness 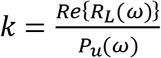 is determined from the real part of the active power spectrum 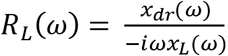, with *x_L_(ω)* and *x_dr_(ω)* as the Fourier transforms of the time series of the positions of the trapping laser and driven bead, respectively, recorded while the trap is oscillating; and the passive power spectrum *P_U_(ω)* = 〈|*x_U_(ω)|^2^*〉, where *x_U_(ω)* is the Fourier transform of the time series of the undriven bead’s thermally fluctuating position while the trap is held stationary.

The generalized Stokes-Einstein relation yields the complex shear modulus as a function of frequency, G*(ω), of each bead’s surrounding microenvironment according to the equation 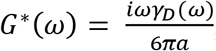 with bead mass *m*, hydrodynamic radius *a*, and the friction relaxation spectrum *γ_D_(ω)*. The complex modulus, *G*\* = |*G*\*|^*(iδ)*^ = *G*’ + *iG*”, has magnitude |*G*|* = *(G’^2^+G”^2^)^1/2^* and loss tangent 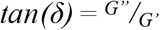 encoding rigidity and hysteresivity, respectively, such that *G’* represents the storage modulus and *G”* represents the loss modulus.

#### Data analysis and statistics

For each cell analyzed, 3-5 beads were measured at different locations within the cell. Only cells exceeding ~30 μm from the coverslip were analyzed. A minimum of 30 cells were analyzed per cell line. Data were analyzed using custom MATLAB and GraphPad Prism programs. Because the modulus magnitudes |*G*\*(*ω*)|, *G’*, and *G”* are log-normal distributed within a single bead, we characterize their central tendency and dispersion, respectively, by the maximum-likelihood estimate of the log-transformed mean 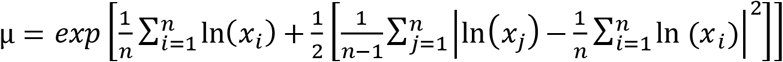 and the maximum-likelihood estimate of the log-transformed variance 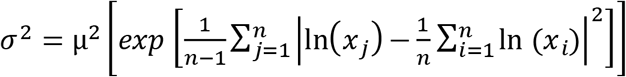. We assume normal distribution of beads within each cell and cells within the cell population, and thus characterize their central tendencies and dispersions respectively by arithmetic mean and variance. Active microrheology data are presented as mean complex, storage, or loss moduli vs frequency from 7 Hz to 15 kHz. The data are also presented as the log 2 transformed pair-wise normalization of the complex modulus, averaged across frequencies (mean ± standard deviation). Data normalized to the complex modulus were analyzed by two-way ANOVA with Tukey’s honestly significant difference post-test.

#### Power Law Fitting

Power law fitting was performed as previously described [16]. Briefly, nonlinear regressions to fit the power law |*G*(ω)*| = *A(ω)^b^* were performed on the mean value at each frequency of |*G*(ω)*| from high frequencies only (400 Hz – 15 kHz). Fits were performed in MATLAB using the Curve Fitting Application, with nonlinear least-squares regression using the Levenberg-Marquardt algorithm and robust weighting with the Least Absolute Residual procedure. The power law exponent *b* is plotted for each individual cell, and *b* values among cell lines were analyzed by one-way ANOVA with Tukey’s multiple comparisons test.

---

**Supplemental Figure 1.**
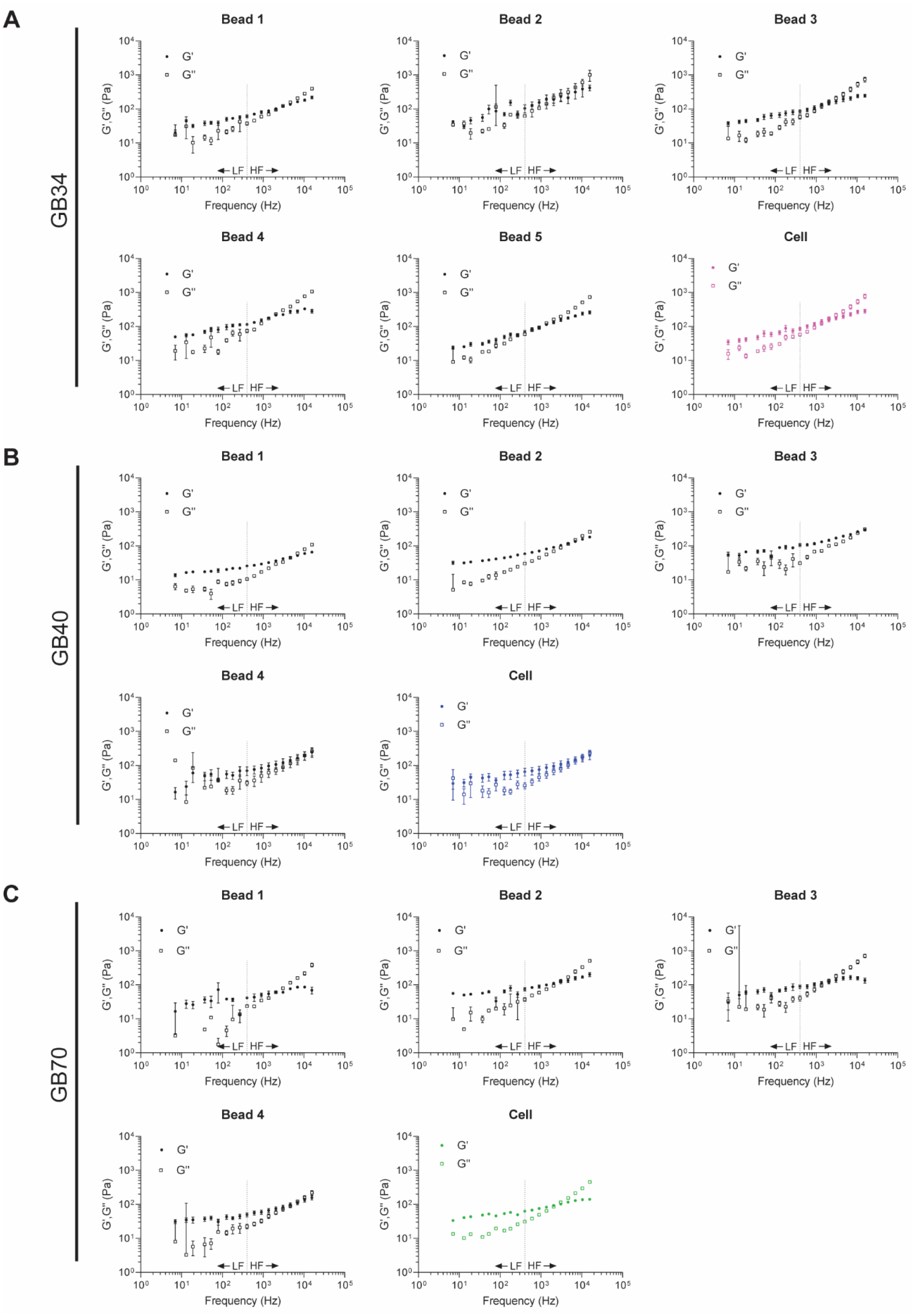
Intracellular mechanical heterogeneity in primary GBM cells. **A-C)** Rheological analysis of representative cells from lines GB34 (**A**), GB40 (**B**), and GB70 (**C**). Bead values for G’ and G” are the average of individual measurements performed on the bead at each frequency (mean ± s.e., n ≤ 7), and cell values are the average (mean ± s.e.) of the beads displayed. Low frequencies (LF) < 400 Hz. High frequencies (HF) >400 Hz.

**Supplemental Figure 2.**
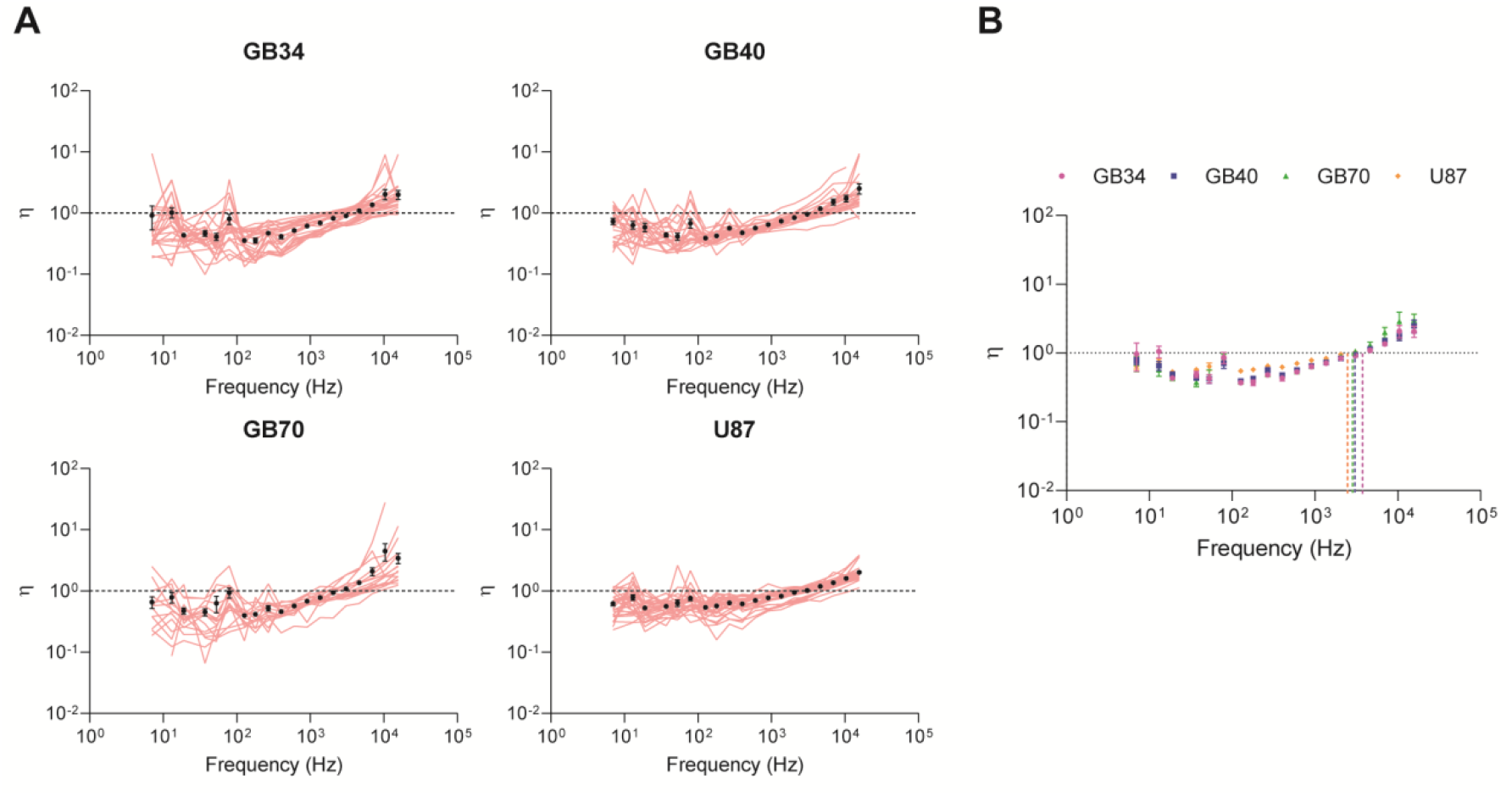
Single cell analysis of hysteresivity (η) among GBM lines. **A)** Mean η(ω) values for each cell within its population, accompanied by the respective population average (mean ± s.e., n ≥ 17 cells). Semifluid materials are described with η=1, while η=0 for purely elastic materials and η is infinite for purely viscous materials. **B)** Population averages for each GBM line (mean ± s.e.). **C)** Pairwise log2 ratios of η(ω) across frequencies comparing all lines. Significance was determined using a 2-way ANOVA (* p < 0.05, ** p < 0.01, *** p < 0.001, **** p < 0.0001).

**Supplemental Figure 3.**
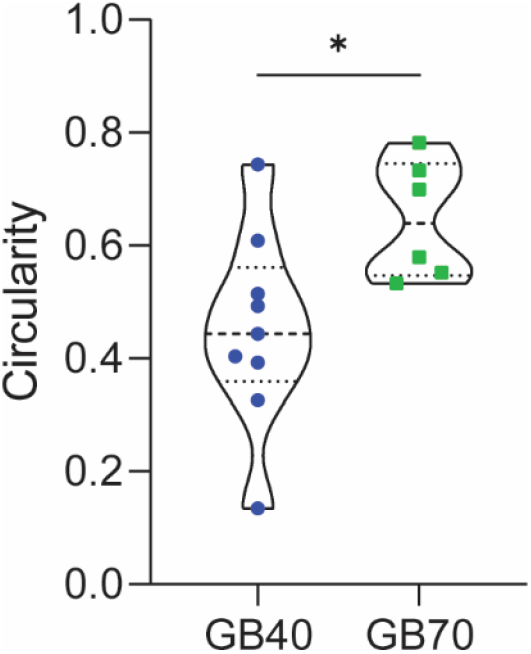
Single cell analysis indicates differences in circularity profiles among primary GBM cell lines. Circularity measurements were performed on micrographs of cells with phalloidin staining of actin. A circularity of 1 indicates a perfect sphere. Measurement was performed in FIJI with built-in circularity plugin according to the formula *circularity = 4π(area/perimeter^2^*). Significance was determined using an unpaired, two-tailed Student’s *t* test (* p < 0.05).

